# LGI1 autoantibodies enhance synaptic transmission by presynaptic K_v_1 loss and increased action potential broadening

**DOI:** 10.1101/2023.10.04.560631

**Authors:** Andreas Ritzau-Jost, Felix Gsell, Josefine Sell, Stefan Sachs, Jacqueline Montanaro, Toni Kirmann, Sebastian Maaß, Sarosh R. Irani, Christian Werner, Christian Geis, Markus Sauer, Ryuichi Shigemoto, Stefan Hallermann

## Abstract

**Background and Objectives:** Autoantibodies against the neuronally secreted protein leucine-rich glioma inactivated 1 (LGI1) cause the most common subtype of autoimmune limbic encephalitis associated with seizures and memory deficits. LGI1 and its receptor ADAM22 are part of a transsynaptic protein complex that includes several proteins involved in presynaptic neurotransmitter release and postsynaptic glutamate sensing. Autoantibodies against LGI1 increase excitatory synaptic strength, but studies that genetically disrupt the LGI1-ADAM22 complex report a reduction in postsynaptic glutamate receptor-mediated responses. Thus, the mechanisms underlying the increased synaptic strength induced by LGI1 autoantibodies remain elusive, and the contributions of presynaptic molecules to the LGI1-transsynaptic complex remain unclear. We therefore investigated the presynaptic mechanisms that mediate autoantibody-induced synaptic strengthening.

**Methods:** We studied the effects of patient-derived purified polyclonal LGI1 autoantibodies on synaptic structure and function by combining direct patch-clamp recordings from presynaptic boutons and somata of hippocampal neurons with super-resolution light and electron microscopy of hippocampal cultures and acute brain slices. We also identified the protein domain mediating the presynaptic effect using domain-specific patient-derived monoclonal antibodies.

**Results:** LGI1 autoantibodies dose-dependently increased short-term depression during high-frequency transmission, consistent with increased release probability. The increased neurotransmission was not related to presynaptic calcium channels, as presynaptic Ca_v_2.1 channel density, calcium current amplitude, and calcium channel gating were unaffected by LGI1 autoantibodies. In contrast, application of LGI1 autoantibodies homogeneously reduced K_v_1.1 and K_v_1.2 channel density on the surface of presynaptic boutons. Direct presynaptic patch-clamp recordings revealed that LGI1 autoantibodies cause a pronounced broadening of the presynaptic action potential. Domain-specific effects of LGI1 autoantibodies were analysed at the soma, where polyclonal LGI1 autoantibodies and patient-derived monoclonal autoantibodies targeting the Epitempin-domain but not the Leucin rich repeat-domain induced action potential broadening.

**Discussion:** Our results indicate that LGI1 autoantibodies do not affect calcium channel density or function, but reduce the density of both K_v_1.1 and K_v_1.2 on presynaptic boutons, thereby broadening the presynaptic action potential and increasing neurotransmitter release. This study provides a molecular explanation for the neuronal hyperactivity induced by LGI1 autoantibodies.

## Introduction

Autoimmune encephalitis is a growing group of diseases caused by autoantibodies against various neuronal antigens, collectively leading to severe mental and behavioral disorders.^1,2^ Limbic encephalitis, primarily affecting the mesial temporal lobe, hippocampus, and amygdala is characterized by focal and generalized seizures and limbic dysfunction including mood changes and amnesia. The most frequent type of autoimmune limbic encephalitis is caused by autoantibodies against the secreted neuronal protein leucine-rich glioma-inactivated 1 (LGI1) resulting in characteristic faciobrachial dystonic and generalized seizures together with mnestic deficits. Seizures rapidly respond to immunotherapy, while patients often develop progressive cognitive impairment and hippocampal sclerosis if treatment is delayed.^3–7^ Besides its major role in limbic encephalitis, mutations in LGI1 have been linked to an inherited form of epilepsy which involves the lateral temporal lobe.^8–10^

LGI1 has two main domains, a N-terminal Leucine-rich repeat (LRR) and a C-terminal epitempin (EPTP) domain.^11^ It is secreted and recruited to the pre- and postsynaptic membrane of excitatory synapses by binding of its EPTP domain to ADAM22-family receptors pre- and postsynaptically.^12–14^ These LGI1-ADAM22 heterodimers have been suggested to dimerize in the synaptic cleft via an LRR-EPTP-interaction, thus linking LGI1-ADAM22s within pre- and postsynaptic membranes to form a *trans*-synaptic tetrameric complex.^13^ ADAM22 receptors have been reported to interact directly or indirectly with both, presynaptic proteins including CAST, SAP97, and various pore-forming or accessory K_v_1 and Ca_v_ channel subunits, and the postsynaptic neurotransmitter receptor scaffold including PSD95 and glutamate receptors.^15,16^ Alternatively, LGI1 was found to be critical for potassium channel expression^17^ and function^18^. The *trans*-synaptic LGI1-ADAM22 complex was therefore proposed as a key component controlling presynaptic transmitter release to postsynaptic receptors.^19^

To study the function of LGI1 at synapses and the consequence of disturbed LGI1 signaling, two main approaches have been adopted. First, genetically modified cell lines or animal models were used that either overexpressed LGI1,^20^ did not express LGI1,^12,21,22^ or that harbored LGI1 mutations associated with inherited epilepsy.^12,20,23,24^ In most of these studies, LGI1 overexpression enhanced and LGI1 loss reduced the postsynaptic α-amino-3-hydroxy-5-methyl-4-isoxazolepropionic acid (AMPA) receptor response and receptor clustering.^12,14,23,25^ Furthermore, LGI1 loss increased the presynaptic neurotransmitter release.^17,20,22^ Second, synaptic LGI1 function was studied using patient-derived polyclonal LGI1 autoantibodies^26–29^ or domain-specific monoclonal autoantibodies^30,31^. Reminiscent of LGI1 loss introduced genetically, incubation with autoantibodies reduced postsynaptic AMPA receptors in primary hippocampal cultures^27^ and acute hippocampal brain slices^28^. Furthermore, recent evidence indicates that LGI1 autoantibodies increase presynaptic release probability and overall synaptic strength^28–30^ with strengthening paralleled by potassium channel loss.^28,31^ However, the mechanism by which LGI1 autoantibodies strengthen presynaptic neurotransmitter release and the responsible molecular domains remain elusive.

Here, we combined electrophysiological somatic and subcellular presynaptic recordings from cultured hippocampal neurons with Stimulated Emission Depletion (STED) microscopy,^32^ Expansion Microscopy (ExM) together with Structured Illumination Microscopy (SIM)^33,34^ and electron microscopy to identify mechanisms involved in the LGI1 autoantibody-mediated increase in presynaptic release. We find that polyclonal LGI1 autoantibodies increase presynaptic release probability independent of calcium channels. In contrast, polyclonal LGI1 autoantibodies reduce presynaptic K_v_1.1 and K_v_1.2 channels and lead to increased action potential broadening, an effect replicated by monoclonal EPTP, but not LRR autoantibodies.

## Results

### LGI1 autoantibodies induce a dose-dependent increase of synaptic release probability

We first investigated the effect of polyclonal LGI1 autoantibodies on excitatory transmission in primary dissociated hippocampal cultures. Cultures were incubated either with patient-derived polyclonal serum LGI1 autoantibodies included in the growth medium for seven days (LGI1-7d; with a second dose applied one day before recordings), with LGI1 autoantibodies for one day only (LGI1-1d), or with patient control IgG antibodies without antineuronal reactivity. We then recorded pharmacologically isolated excitatory postsynaptic currents (EPSCs) in somatic whole-cell voltage-clamp recordings evoked by external stimulation. EPSCs in control IgG-incubated neurons showed facilitation at frequencies of 20 Hz and 50 Hz, reflected in paired pulse ratios (PPRs) > 1. Incubation with LGI1 autoantibodies reduced PPRs at both frequencies in a dose-dependent manner, indicating that LGI1 autoantibodies increase the synaptic release probability (Fig. 1A and B; for example, median [IQR] PPR at 20 Hz: 1.03 [0.84–1.24] and 0.67 [0.53–0.69], n = 16 and 11, for control and LGI1-7d, respectively; non-parametric Kruskal-Wallis ANOVA test P < 0.001 and post-hoc test P < 0.001 for control and LGI1-7d). To study autoantibody-induced changes in synaptic transmission in more detail, we analyzed short-term plasticity during evoked EPSC trains (50 EPSCs at 20 Hz; Fig. 1C). LGI1 autoantibody-incubation suppressed facilitation and induced faster and stronger depression of excitatory currents, in line with higher presynaptic release probability upon autoantibody incubation (Fig. 1C; median [IQR] amplitude of the 10 last train EPSCs normalized to the 1^st^ train EPSC: 0.42 [0.40–0.56] and 0.28 [0.24–0.29], n = 14 and 11, for control and LGI1-7d, respectively, post-hoc P = 0.004). These data indicate a dose dependent increase in presynaptic release probability upon treatment with LGI1 autoantibodies. Because seven-day antibody-treatment more robustly affected synaptic transmission than one day-treatment, we adopted the seven day-treatment for subsequent analyses.

**Figure 1.**
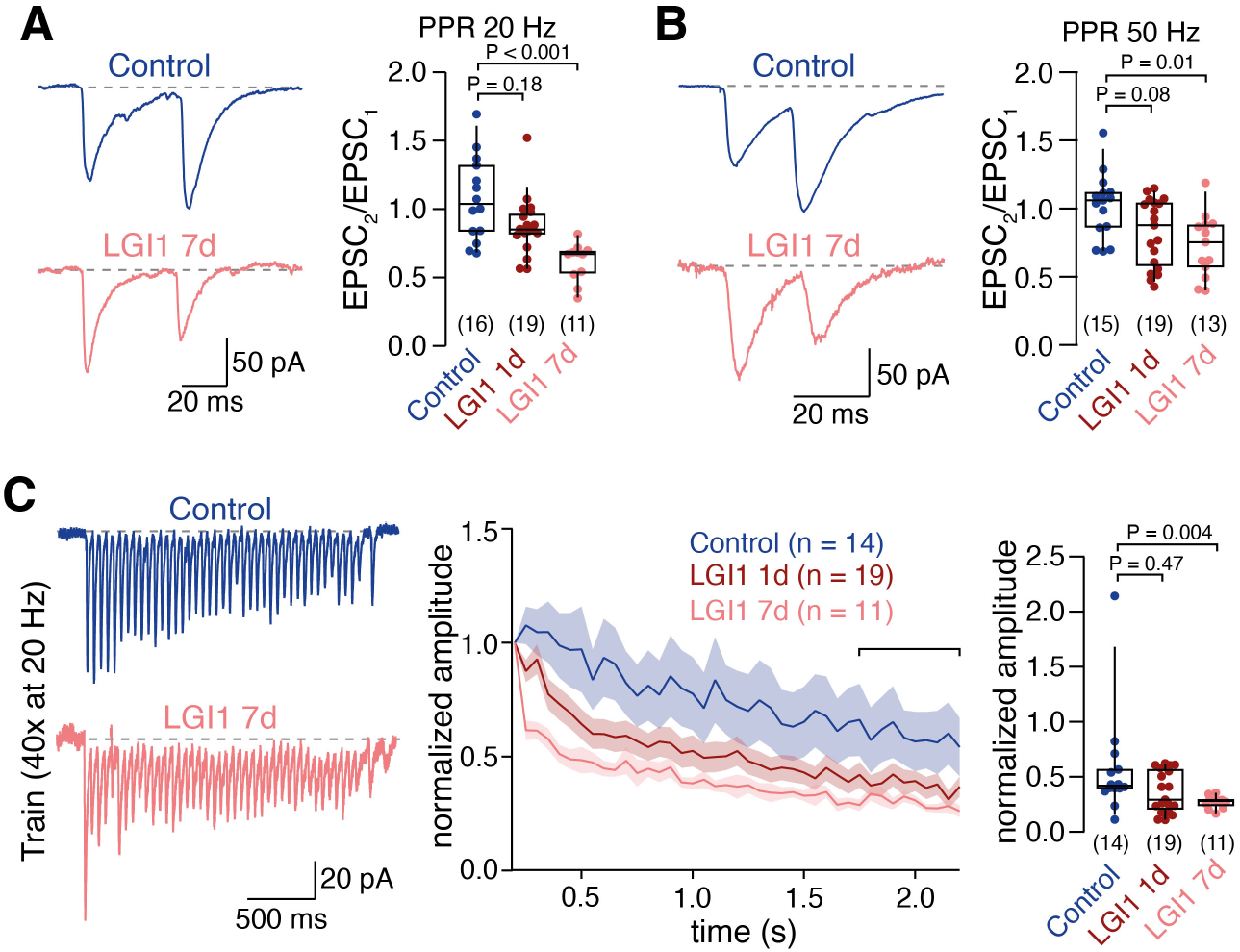
LGI1 autoantibodies induce a dose-dependent increase of synaptic release probability. **(A)** *Left:* Example paired EPSCs evoked at 20 Hz under control condition (blue) and following 7 days of LGI1 autoantibody incubation (orange). *Right:* Paired pulse ratio (EPSC_2_/EPSC_1_) at 20 Hz under control condition (blue) and following 1 day or 7 days of LGI1 autoantibody incubation (red and orange, respectively). A non-parametric ANOVA (Kruskal-Wallis) test revealed P < 0.001. **(B)** Same data as in A but for 50 Hz. A non-parametric ANOVA (Kruskal-Wallis) test revealed P = 0.012. **(C)** *Left:* Example EPSC train (40 EPSCs evoked at 20 Hz) under control condition (blue) and following 7 days of LGI1 autoantibody incubation (orange). *Middle:* Normalized train EPSC amplitudes in control condition (blue) and following 1 day or 7 days of LGI1 autoantibody incubation (red and orange, respectively; mean ± SEM). *Right:* Depression of normalized EPSC amplitudes during the late phase of the train (see bracket in middle panel). A non-parametric ANOVA (Kruskal-Wallis) test revealed P = 0.035 Numbers in brackets reflect recordings from individual neurons. Box plots cover percentile 25–75 with median indicated, whiskers indicate percentiles 10–90. The P values of the non-parametric ANOVA (Kruskal-Wallis) tests are provided in the legends and the P-values of the non-parametric post-hoc tests (Dwass-Steel-Critchlow-Fligner pairwise comparisons) are provided in the figures.

### LGI1 autoantibodies have little effect on presynaptic Cav2.1 calcium channel density

The release probability for synaptic vesicles is influenced by the number, the position, and the properties of the presynaptic calcium channels.^35–37^ Furthermore, proteome studies indicate interactions between the LGI1-receptor ADAM22 and calcium channels.^19,38^ Therefore, a straightforward explanation for the autoantibody-induced increase in release probability could be an increased presynaptic calcium influx due to either higher calcium channel abundance or faster channel gating. To first test whether LGI1 autoantibodies affected presynaptic calcium channel abundance, we performed STED imaging of Ca_v_2.1 channels, which is one of the main calcium channel types at hippocampal synapses.^39,40^ Ca_v_2.1 fluorescence signal intensities were quantified at excitatory presynapses (labeled by the vesicular glutamate transporter vGlut1) and excitatory active zones (labeled by Bassoon within vGlut1-postive presynapses). LGI1 autoantibodies decreased Ca_v_2.1 channel fluorescence intensities within both, presynapses and excitatory active zones (Fig. 2A, B; median [IQR] Ca_v_2.1 intensity within vGlut1-positive Bassoon: 18.7 [25.8–12.9] and 16.3 [22.6–11.4], n = 2949 and 2857 synapses for control and LGI1, respectively, P < 0.001; on average 10.1% less Ca_v_2.1 within Bassoon in 4 cultures). This ∼10% decrease in Ca_v_2.1 channels by LGI1 autoantibodies is in contrast to an increased synaptic release probability, which would require increased presynaptic Ca_v_2.1 channels instead. To analyze the effect of LGI1 autoantibodies on Ca_v_2.1 channels within presynaptic active zones in more detail, we performed freeze-fracture replica immunoelectron microscopy of hippocampal presynapses from mice chronically infused with LGI1 autoantibodies using intraventricular osmotic pumps. We first analyzed dentate gyrus perforant path–granule cell synapses, as LGI1 expression is highest in the dentate gyrus^18^ and transmission is affected presynaptically upon genetic alteration of LGI1^20^ or LGI1 autoantibodies^28^. Chronic LGI1 autoantibody-infusion did not affect active zone Ca_v_2.1 channel density (Fig. 2C; median [IQR] Ca_v_2.1 particle density per µm^2^: 248 [186–444] and 220 [156–386], 96 and 117 synapses for control and LGI1, respectively, from 3 animals each, P = 0.12) with a trend towards a ∼10% reduction upon autoantibody-treatment, similar to STED recordings in cultured neurons. Additionally, we quantified Ca_v_2.1 channel density at another LGI1-expressing synapse between dentate mossy fibers and CA3 neurons from chronically infused mice. Similarly, LGI1 autoantibodies did not affect Ca_v_2.1 channel density at these synapses (Fig. 2D; median [IQR] Ca_v_2.1 particle density per µm^2^: 152 [100–225] and 147 [91–213], 234 and 240 synapses for control and LGI1, respectively, from 6 animals each, P = 0.18). These data show that LGI1 autoantibodies caused, if anything, a small reduction in presynaptic Ca_v_2.1 channel density in boutons of hippocampal cultures and in tissue. Therefore, alterations in calcium channels density cannot explain increased release probability by LGI1 autoantibodies.

**Figure 2.**
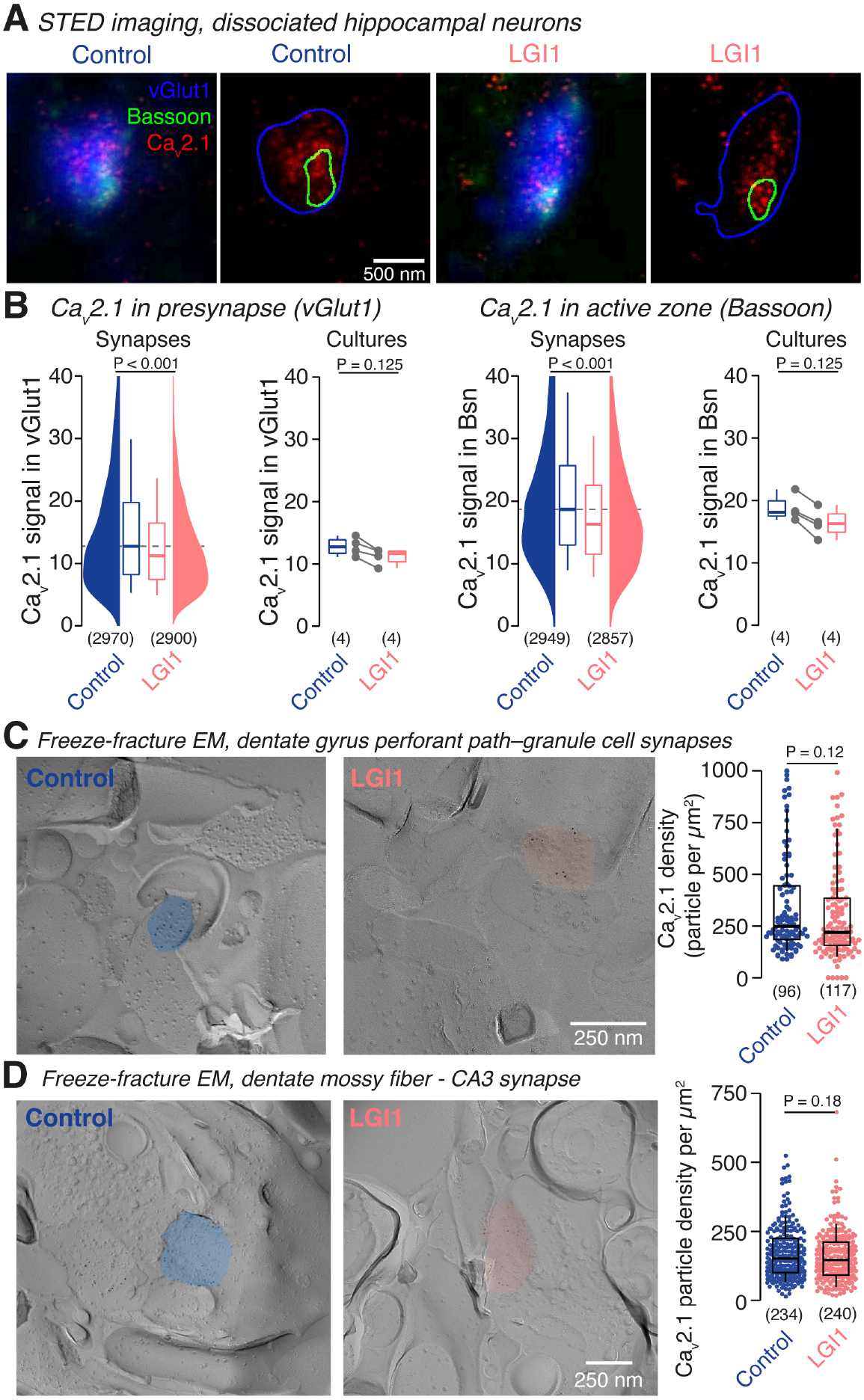
LGI1 autoantibodies have little effect on presynaptic Ca_v_2.1 calcium channel density. **(A)** STED fluorescence images of synapses stained for vGlut1 as a synaptic marker (blue), Bassoon as an active zone marker (green), and Ca_v_2.1 channels (red) in control cultures (*left*) and cultures treated with LGI1 autoantibodies (*right*). Next to fluorescence images, images following segmentation are provided showing the boundaries of the marker channels. **(B)** Presynaptic Ca_v_2.1 fluorescence (within vGlut1, *left*) and active zone Ca_v_2.1 fluorescence (within Bassoon, *right*) in control synapses (*blue*) and LGI1 autoantibody-treated synapses (*orange*) for all recorded synapses (‘Synapses’) and across all recorded cultures (‘Cultures’). **(C)** Electron microscopic images of freeze-fracture replica immunolabeling for Ca_v_2.1 in hippocampal perforant path-granule cell synapses for control and following LGI1 autoantibody-treatment (color code as in B). Ca_v_2.1 particle densities in synapses pooled from 3 animals per group. **(D)** Electron microscopic images of freeze-fracture replica immunolabeling for Ca_v_2.1 in hippocampal dentate mossy fiber-CA3 synapses for control and following LGI1 autoantibody-treatment (color code as in B). Ca_v_2.1 particle densities in synapses pooled from 6 animals per group. Numbers in brackets reflect individual presynapses (B–D) or cultures (B). Box plots cover percentile 25–75 with median indicated, whiskers indicate percentiles 10–90. P-values were calculated using the Mann–Whitney *U* test.

### LGI1 autoantibodies do not affect presynaptic calcium channel gating

The LGI1-receptor ADAM22 has been shown to interact with various Ca_v_ channel beta-subunits ^19^, which in turn affect calcium current gating ^41^. We therefore determined the effect of LGI1 autoantibodies on calcium channel gating by directly measuring pharmacologically-isolated presynaptic calcium currents in whole-cell voltage-clamp recordings from boutons in hippocampal cultures (Fig. 3A). Calcium currents upon 3-ms depolarization were similar in amplitudes for control and LGI1 autoantibody-treated boutons (Fig. 3B; median [IQR] current amplitude at 0 mV: 18.1 [10.5–28.4] pA and 18.3 [10.5–25.3] pA, n = 10 and 17, for control and LGI1, respectively, P = 0.66; 2-way-ANOVA for overall effect: P = 0.079). In addition, the time course of current activation was not affected by LGI1 autoantibodies (Fig. 3C; median [IQR] activation time constant at 0 mV: 0.73 [0.54–1.03] ms and 0.66 [0.46–1.25] ms, n = 13 and 20 for control and LGI1, respectively, P = 0.86). Similarly, amplitude and time course of calcium current inactivation were unchanged following LGI1 autoantibody-treatment (Fig. 3D, E; median [IQR] current amplitude at, e.g., –30 mV: 7.9 [1.9–45.7] pA and 7.8 [2.44–14.5] pA, n = 11 and 13 for control and LGI1, respectively, P = 0.80; 2-way-ANOVA for overall LGI1 autoantibody effect: P = 0.18; median [IQR] inactivation time constant at –30 mV: 0.49 [0.25–1.09] ms and 0.60 [0.35–1.01] ms, n = 11 and 15 for control and LGI1, respectively, P = 0.78; 2-way-ANOVA for overall LGI1 autoantibody effect: P = 0.78). Unaltered presynaptic calcium currents indicate that LGI1 autoantibodies did not affect calcium channel gating and thus cannot explain the increased release probability by LGI1 autoantibodies.

**Figure 3.**
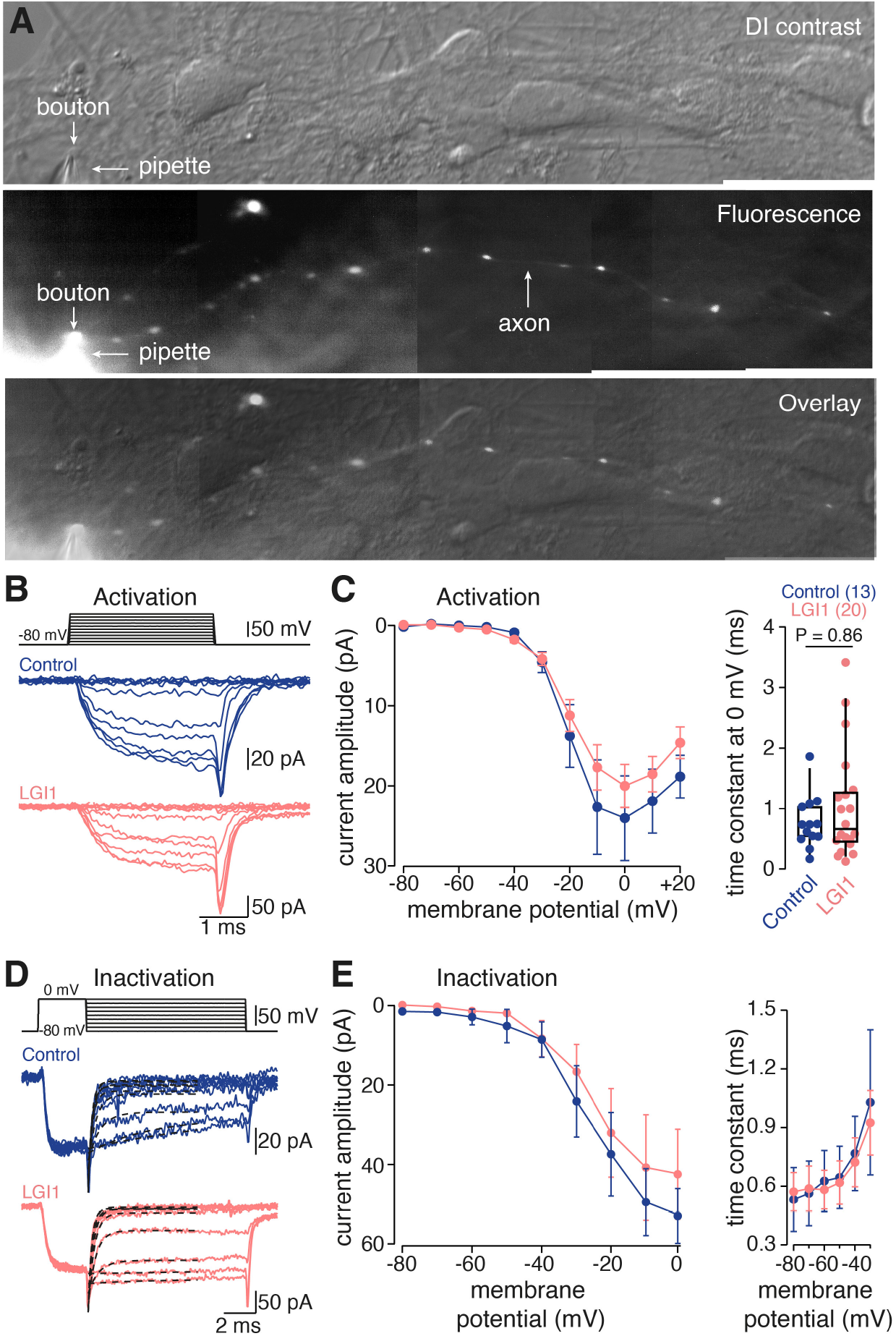
LGI1 autoantibodies do not affect presynaptic calcium channel gating. **(A)** Difference-interference contrast (DIC) image, fluorescence image (200 µM Atto 488 contained in recording pipette), and an overlay of both of a small bouton whole-cell recording in primary hippocampal cultures. **(B)** Pharmacologically-isolated calcium currents evoked by 3 ms-depolarizations to different voltages in control (blue) and LGI1 autoantibody-treated boutons (orange). **(C)** Calcium current amplitudes during the final 1 ms of the step depolarization (*left*, mean ± SEM) and activation time constant at 0 mV (*right*) in control and LGI1 autoantibody-treated boutons (color code as in B). **(D)** Calcium currents evoked with an inactivation paradigm (3 ms depolarization to 0 mV followed by a 10 ms inactivation step at different voltages) in control and LGI1 autoantibody-treated boutons (color code as in B). **(E)** Calcium current amplitudes during the final 1 ms of inactivation steps to different voltages as in D (*left*) and inactivation time constant at various potentials (*right*) in control and LGI1 autoantibody-treated boutons (color code as in B; mean ± SEM). Numbers in brackets reflect recordings from individual presynaptic boutons. Box plots cover percentile 25–75 with median indicated, whiskers indicate percentiles 10–90. The P-value was calculated using the Mann–Whitney *U* test (*right* in C).

### Nanoscale localization of presynaptic Kv1.1 and Kv1.2 channels in hippocampal synapses

LGI1 interacts via ADAM22-receptors with K_v_1 potassium channels ^18,19^, which are reduced in animals treated with LGI1 autoantibodies ^28,31^. We therefore hypothesized that the release probability is increased because of the loss of presynaptic K_v_1 channels. We first tested whether K_v_1.1 and K_v_1.2 channel subtypes, which have been linked to LGI1 functionally and in biochemical essays ^12,18,19^, are localized presynaptically in cultured hippocampal neurons. Using structured illumination microscopy (SIM) of neurons co-stained for Bassoon, we found that K_v_1.2 channels localized more to the presynaptic active zone in cultured hippocampal neurons compared to K_v_1.1 (Fig. 4A; Mander’s colocalization coefficients for K_v_1.1 and Bassoon = 0.28 ± 0.11 (mean ± SD), n = 137 synapses; K_v_1.2 and Bassoon = 0.62 ± 0.13, n = 195 synapses; the mean per each of the three cultures were 0.26, 0.27, and 0.30 for K_v_1.1 and 0.66, 0.62, and 0.58 for K_v_1.2). To test potential limitations of the spatial resolution, we used post-gelation immunolabeling ExM in combination with SIM (Ex-SIM).^34^ Samples expanded ∼7.5-fold, thus enabling a spatial resolution of ∼20 nm by multicolor SIM. The data indicate that both K_v_1.1 and K_v_1.2 channels localized at vGlut1-positive presynaptic nerve terminals of cultured hippocampal neurons. Furthermore, both K_v_1.1 and K_v_1.2 were found inside and outside of the Bassoon-labeled active zone in cultured hippocampal neurons while K_v_1.2 seemed more clustered at the active zone. To corroborate the presence of K_v_1 channels at hippocampal presynapses in brain tissue, we studied hippocampal dentate gyrus perforant path–granule cell synapses in perfusion-fixed hippocampal tissues. At these synapses, it was previously shown that LGI1 antibodies also increase the release probability ^28^. Using pre-embedding electron microscopy, we localized K_v_1.1 and K_v_1.2 channels at the perforant path– granule cell synapses (Fig. 4C). A three-dimensional reconstruction of perforant path axon terminals and the adjacent axons indicated a homogeneous K_v_1.1 channel distribution (Fig. 4D), consistent with <5% of both channel subtypes localized at or close to the small surface area building the presynaptic active zone (*active zone* and *perisynaptic*; Fig. 4D). However, the large majority of the K_v_1.1 and K_v_1.2 channels were found outside of the active zone. Thus, several complementary high-resolution light and electron microscopic techniques confirm the localization of both K_v_1.1 and K_v_1.2 channels at hippocampal presynapses and indicate a rather homogeneous distribution.

**Figure 4.**
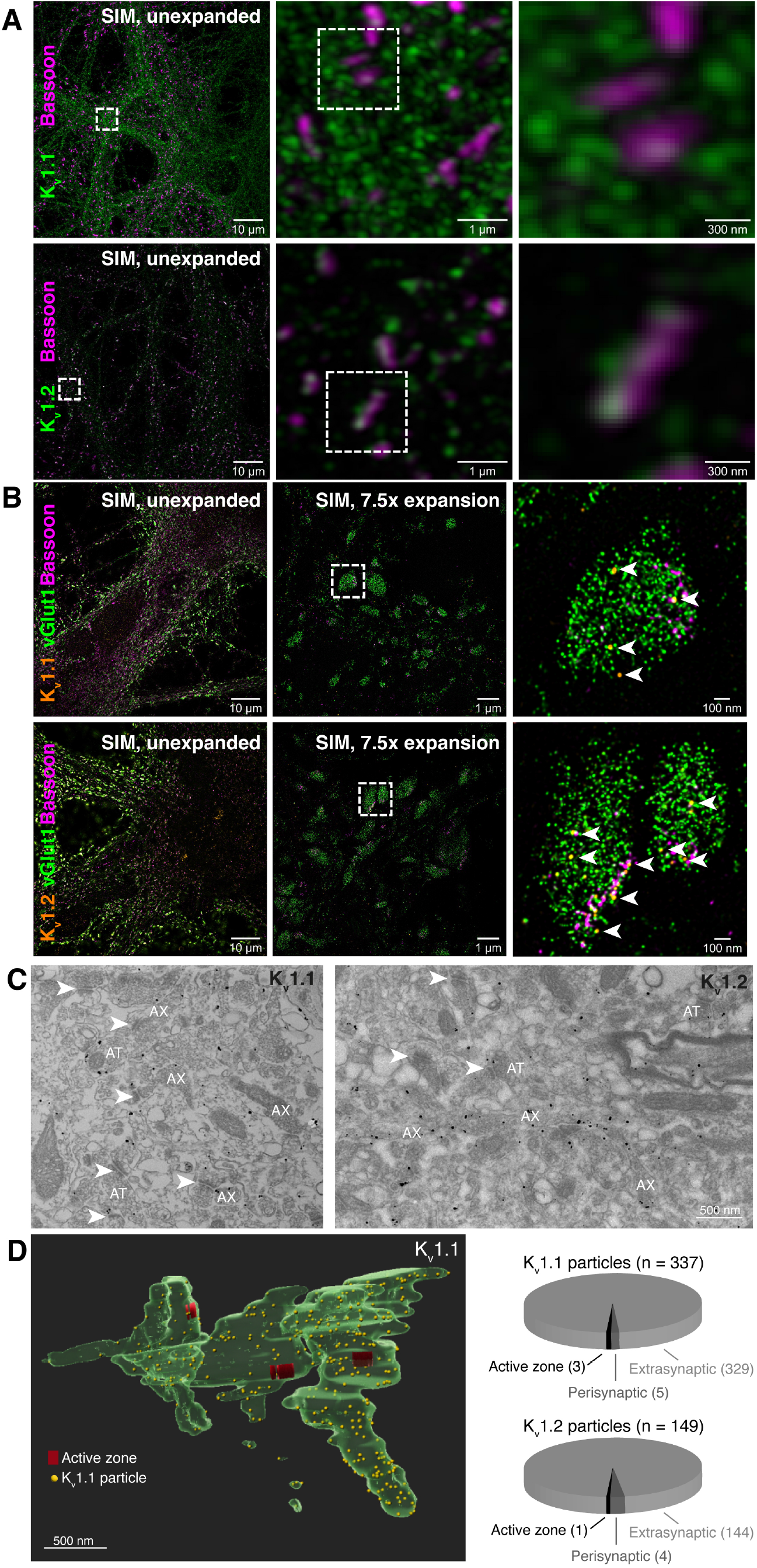
Nano-scale presynaptic localization of K_v_1.1 and K_v_1.2 channels in hippocampal synapses. **(A)** Structured illumination microscopic (SIM) images of synapses in primary hippocampal cultures labeled for Bassoon and K_v_1.1 (*top*) or K_v_1.2 channels (*bottom*) at increasing magnification (boxed areas, from left to right). **(B)** SIM images of unexpanded cultures and cultures expanded ∼7.5-fold, triple-stained for vGlut, Bassoon and K_v_1.1 (*top*) or K_v_1.2 channels (*bottom*). Right images are magnifications of expanded SIM-images (boxed area), arrow heads indicate K_v_1 clusters. **(C)** Electron microscopic image of a dentate gyrus molecular layer section immunogold-labelled for K_v_1.1 (*left*) or K_v_1.2 channels (*right*; AX = axon, AT = axon terminal; arrowheads depict excitatory synapses with postsynaptic electron densities) **(D)** 3D reconstruction of axons and their terminals (*left*) harboring multiple active zones (red) and K_v_1.1 particles (yellow spheres) and quantification of K_v_1.1 and K_v_1.2 particle localization (*right*) within active zones (black), the perisynaptic space (dark grey; ≤ 60 nm from the active zone edge) and the extrasynaptic space (light grey; > 60 nm from the active zone edge). Numbers in brackets indicate total particle counts and counts within respective localizations.

### LGI1 autoantibodies reduce presynaptic Kv1.1 and Kv1.2 channels

After we found presynaptic localization of K_v_1.1 and K_v_1.2 channels, we tested whether their localization was affected by LGI1 autoantibodies. LGI1 autoantibodies were previously shown to reduce general synaptic K_v_1.1 channels using confocal imaging^28^ or western blots^31^. We performed STED imaging due to higher resolution compared to confocal microscopy and higher throughput compared to Ex-SIM technique and investigated K_v_1.1 and K_v_1.2 channels specifically at excitatory presynapses (Fig. 5A). Because of the variability in K_v_1 signal intensity between synapses, we repeated the antibody application, immunostaining, image acquisition, and image analyses in 10 cultures independently generated from 10 different animals. When we merged all analyzed synapses, LGI1 autoantibodies reduced K_v_1.1 and K_v_1.2 signals at excitatory presynapses (Fig. 5B, D) and active zones (Fig. 5C, E) by 10–15% (*Synapses* in Fig. 5B–E; median change for K_v_1.1 within vGlut1 –13.6%, K_v_1.2 within vGlut1 –13.2%, K_v_1.1 within Bassoon –12.3%, K_v_1.2 within Bassoon –10.2%, each P < 0.001 and n = ∼5000 synapses). Even when we analyzed each culture separately (which might represent an over-critical definition of the biological replicate), the LGI1 autoantibodies showed trends of reduction or statistically significant reduction of both K_v_1.1 and K_v_1.2 within vGlut1 and Bassoon (*Cultures* in Fig. 5B–E). Consistent with a homogeneous distribution of K_v_1.1 and K_v_1.2 in electron microscopy, we also observed a reduction in the density of presynaptic K_v_1.1 and K_v_1.2 outside of the active zone (i.e., inside the vGlut1 but outside of the Bassoon mask; data not shown). Thus, presynaptic K_v_1.1 and K_v_1.2 channels are both homogeneously reduced upon treatment with LGI1 autoantibodies.

**Figure 5.**
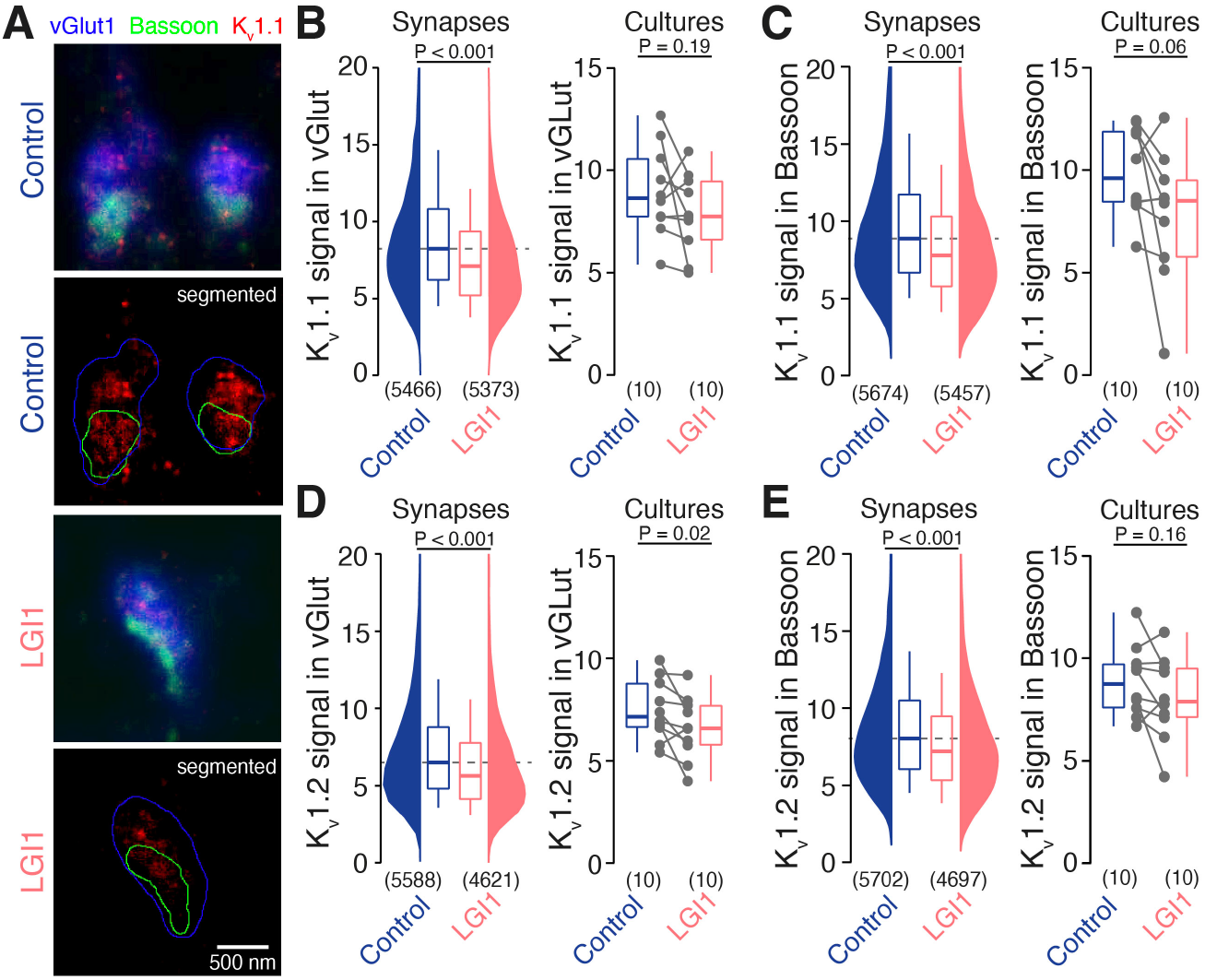
LGI1 autoantibodies reduce presynaptic K_v_1.1 and K_v_1.2 channels. **(A)** Fluorescence images of primary hippocampal cultures stained for vGlut1 (blue), Bassoon (green), and K_v_1.1 (red) following treatment with control (blue, upper panels) or LGI1 autoantibodies (orange, lower panels). Segmented vGlut1 and Bassoon masks are provided below the unsegmented images. **(B)** K_v_1.1 fluorescence intensity within vGlut1 masks for control (blue) and LGI1 autoantibody-treatment (orange) across all recorded synapses (*left*) or cultures (*right*). **(C)** K_v_1.1 fluorescence intensity within Bassoon masks of control and LGI1 autoantibody-treated synapses (*left*) or cultures (*right*), color code as in B. **(D)** K_v_1.2 fluorescence intensity within vGlut1 masks of control and LGI1 autoantibody-treated synapses (*left*) or cultures (*right*), color code as in B. **(E)** K_v_1.2 fluorescence intensity within Bassoon masks of control and LGI1 autoantibody-treated synapses (*left*) or cultures (*right*), color code as in B. Box plots provide median and cover percentile 25–75, whiskers reflect percentiles 10–90. Broken lines indicate the respective control condition median intensity. Numbers in brackets provide number of total analyzed synapse or independent cultures. P-values were calculated using the Mann–Whitney *U* test.

### LGI1 autoantibodies lead to increased presynaptic AP broadening

K_v_1 channels control presynaptic action potential duration (e.g., see ref. 42 and references therein). To determine the functional relevance of presynaptic K_v_1 channel loss, we performed direct current-clamp recordings from boutons in dissociated hippocampal cultures^42,43^ following autoantibody-treatment. Action potentials evoked by current-injections had large amplitudes and short half-durations (quantified as full-width recorded at half-maximal amplitude, FWHM), similar to previous findings at boutons of dissociated neocortical cultures.^42^ Changes in action potential shape were tested by evoking trains of 90 action potentials at 20 or 50 Hz (Fig. 6A). Action potential broadening during train stimulation was pronounced following LGI1 autoantibody-treatment (Fig. 6B and C; 20 Hz: median [IQR] broadening of the last 10 APs: 29.1 [25.0–37.7] % and 48.4 [25.6–77.1] %, n = 17 and 15 for control and LGI1, respectively, P = 0.05; 50 Hz: 46.9 [38.0–67.2] % and 69.9 [40.8–173.9] %, n = 16 and 16 for control and LGI1, respectively, P = 0.03). Due to the large bouton-to-bouton variability, the absolute duration of the last 10 AP only showed a trend towards an increase duration (P = 0.23 and P = 0.16 for 20 and 50 Hz, respectively; data not shown). In contrast to the duration of action potentials, the amplitudes of presynaptic action potentials were not affected by LGI1 autoantibodies (eFigure 1; change in median amplitude of last 10 APs < 3% at 20 Hz and < 10% at 50 Hz, both P > 0.05). Besides changes in action potential broadening, incubation with LGI1 autoantibodies also increased bouton excitability, leading to aberrant action potential firing during current injections (Fig. 6D; repetitive action potentials upon prolonged current injections in 1/15 and 5/13 boutons for control and LGI1, respectively, P = 0.04). Similar to presynaptic action potentials, somatic action potentials were broadened and showed increased activity-induced broadening following incubation with LGI1 autoantibodies (eFigure 2). It is well established that broader presynaptic action potentials lead to more calcium influx and higher release probability.^44–48^ Thus, these data indicate that, by reducing K_v_1 channels, LGI1 autoantibodies enhance somatic and presynaptic action potential broadening and thus synaptic release probability.

**Figure 6.**
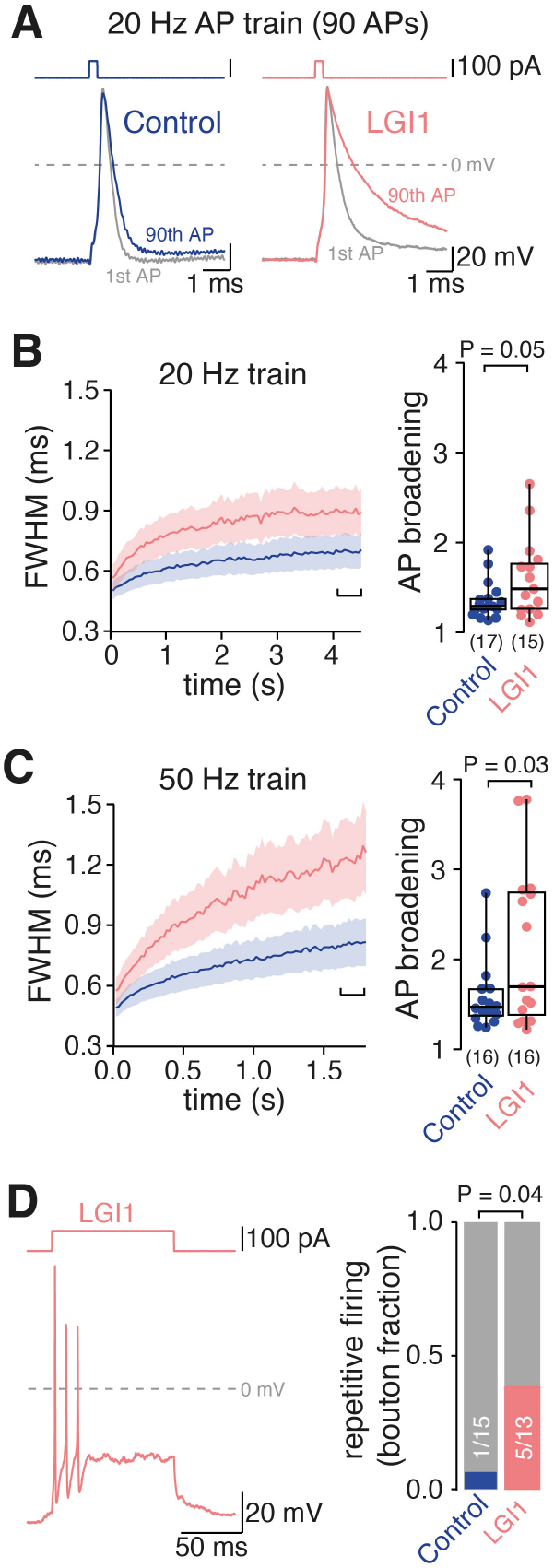
LGI1 autoantibodies lead to increased presynaptic AP broadening. **(A)** Overlay of first and last action potential of an action potential train (90 APs at 20 Hz) following incubation with control (blue) or LGI1 autoantibodies (orange). **(B)** Time course (*left*; mean ± SEM) and magnitude of action potential broadening (*right*; AP broadening; FWHM of 10 last train action potentials normalized to the first action potential) during 20 Hz train stimulation in control and LGI1 autoantibody-treated presynaptic boutons (blue and orange, respectively). **(C)** Action potential broadening as in C for 50 Hz train stimulation. **(D)** Example recording and fraction of boutons showing multiple action potentials upon prolonged current injection in control and LGI1 autoantibody-treated boutons. Box plots provide median and cover percentile 25–75, whiskers reflect percentiles 10–90. Numbers in brackets provide number of recorded presynaptic boutons. The P-values were calculated using the Mann–Whitney *U* test (B, C) or the Chi-square test (D).

### Only autoantibodies against the EPTP-domain of LGI1 cause action potential broadening

To address which of the two main LGI1 domains is involved in the antibody-mediated action potential broadening, we again recorded somatic action potentials, this time however following treatment with patient-derived monoclonal autoantibodies specifically targeting either only the EPTP or the LRR domain.^31^ LRR autoantibodies did not affect action potential broadening during 20 or 50 Hz trains compared to cells treated with control isotype monoclonal antibodies (Fig. 7A–C; median [IQR] FWHM of last 10 train APs at 20 Hz: 1.08 [0.98–1.20] ms and 1.20 [0.99–1.31] ms, n = 17 and 22 for control and LLR, respectively, P = 0.45; 50 Hz:1.42 [1.28–1.46] ms and 1.39 [1.14–1.58] ms, n = 15 and 12 for control and LLR, respectively, P = 0.82). In contrast, treatment with EPTP autoantibodies led to enhanced action potential broadening during 20 and 50 Hz trains (Fig. 7A–C; median [IQR] FWHM of last 10 train APs at 20 Hz: 1.32 [1.12–1.46] ms for EPTP, n = 17, P = 0.027; 50 Hz: 1.90 [1.55–2.10] ms for EPTP, n = 9, P = 0.023). The broadening of somatic action potentials following EPTP autoantibody treatment was similar in magnitude to broadening of the somatic action potential induced by polyclonal LGI1 autoantibodies (cf. Fig. 7B and C and eFigure 2B and C), indicating that antibody binding to the EPTP domain underlies action potential broadening.

**Figure 7.**
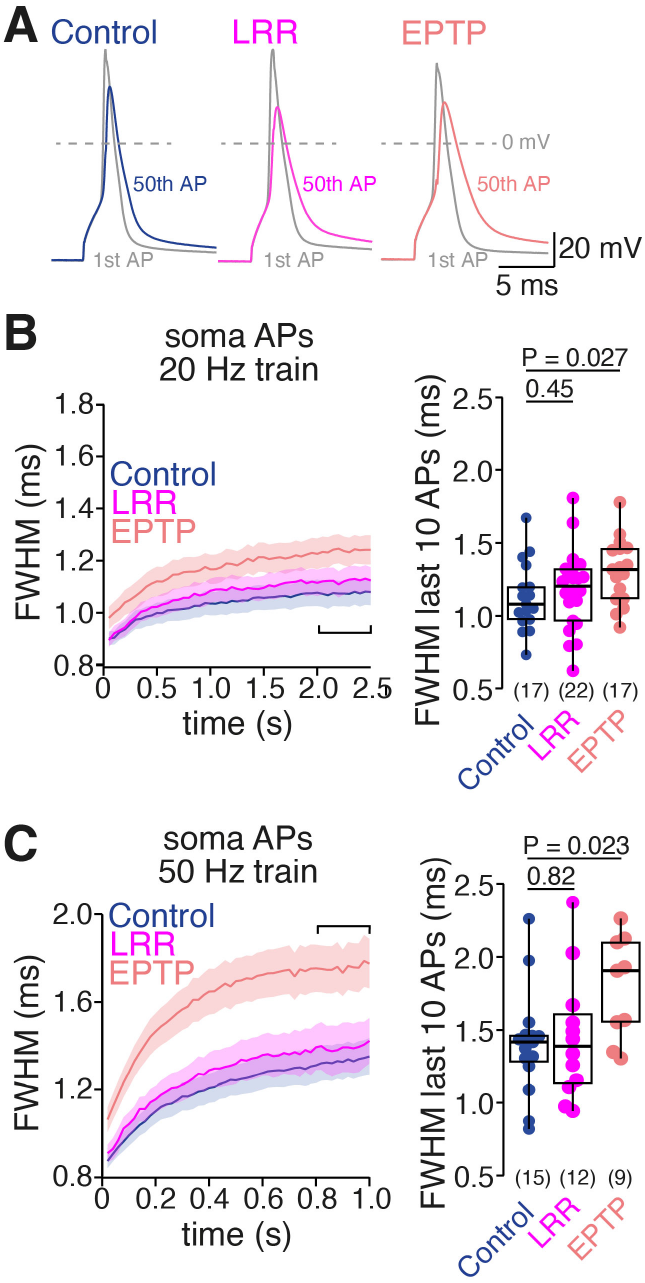
Only autoantibodies against the EPTP-domain of LGI1 cause action potential broadening. **(A)** Overlay of first and last action potentials of a somatic action potential train (50x at 20 Hz) recorded after treatment with control antibodies (blue), LRR autoantibodies (magenta), or EPTP autoantibodies (orange). **(B)** Time course (*left*; mean ± SEM) and magnitude of somatic action potential broadening (*right*; FWHM of 10 last train action potentials) during 20 Hz action potential trains following treatment with control antibodies, LRR autoantibodies, or EPTP autoantibodies (color code as in A) **(C)** Somatic action potential broadening as in B for 50 Hz train stimulation. Box plots provide median and cover percentile 25–75, whiskers reflect percentiles 10–90. Numbers in brackets provide number of recorded presynaptic boutons. P-values were calculated using the using the Kruskal-Wallis tests followed by Dunn’s multiple comparison tests.

## Discussion

Our results have important implication for understanding the pathophysiology of LGI1 autoimmune encephalitis and the physiological functions of LGI1. In particular, our study demonstrates that: (1) LGI1 autoantibodies broaden presynaptic action potentials, which explains the observed increase in release probability. (2) We did not find relevant changes in the density nor the gating of calcium channels upon LGI1 autoantibody treatment. (3) The homogeneous presynaptic distribution and reduction of K_v_1.1 and K_v_1.2 channels upon LGI1 autoantibody treatment indicate that LGI1 can act outside of the release site in addition to its transsynaptic function. (4) Experiments with domain-specific patient-derived monoclonal autoantibodies indicate that action potential broadening is mediated by autoantibodies targeting the EPTP domain, but not by antibodies targeting the LRR domain. Thus, our study provides a mechanistic framework explaining the neuronal hyperactivity of patients with LGI1 antibody encephalitis.

### Presynaptic effects of LGI1 autoantibodies on synaptic transmission

It is controversial whether LGI1 autoantibodies affect synaptic transmission presynaptically, postsynaptically, or both, pre- and postsynaptically. We found that LGI1 autoantibodies decreased paired-pulse ratios and increased synaptic depression arguing for a presynaptic effect of LGI1 autoantibodies.^49^ Our results are consistent with previous studies reporting that LGI1 autoantibodies increase synaptic strength and decrease paired-pulse ratio at hippocampal perforant path-granule cell synapses and reduce synaptic failures in CA1 neurons^28^, with a similar trend towards higher mEPSC frequency observed in CA3 neurons.^30^ Synapses onto hippocampal CA1 and CA3 neurons were not affected in strength or paired-pulse ratio by LGI1 autoantibodies ^26,28,29,31^, which might be due to lower abundance of the LGI1 protein at these synapses.^18^ The increased release probability upon LGI1 antagonism provides an explanation for the hyperactivity in both LGI1 autoantibody-treated neurons^26,29^ and neurons of LGI1 knock-out mice^12,21,22,50^. Furthermore, the increased release probability might also serve as a basis for the epileptic seizures of patients suffering from LGI1 antibody encephalitis.^51^ The faciobrachial dystonic seizures respond intriguingly fast to immunotherapy, whereas antiseizure medication is often ineffective.^5^ Our data suggest that the ineffectiveness of antiseizure medication could be due to the increased presynaptic function. However, in an observational study levetiracetam (interfering with glutamate release) appears to be less effective compared to carbamazepine (interfering with sodium channels),^52^ which argues against this assumption. More studies are needed to better understand the cause and treatment of seizures in anti-LGI1 encephalitis.

### Molecular mechanisms for the LGI1 autoantibody-mediated increase in release probability

LGI1 and ADAM receptor proteins have previously been shown to affect K_v_1 channel gating^18^ and expression^17,53^, and LGI1 autoantibodies immunoprecipitated with K_v_ channels^4,16^. We found that both K_v_1.1 and K_v_1.2 subunits were localized presynaptically, consistent with previous results on K_v_1 channel localization.^54^ LGI1 autoantibodies reduced presynaptic K_v_1.1 and K_v_1.2 channels, in agreement with reduced hippocampal K_v_1.1 fluorescence^28^ and K_v_1 protein levels^31^ following autoantibody treatment. Direct bouton patch-clamp recordings revealed enhanced action potential broadening during train stimulation, a well-known consequence of reduced K_v_1 conductance upon activity-dependent K_v_1 channel inactivation^44,45^ or pharmacological K_v_1 channel block (e.g., see ref. 42 and references therein). Therefore, our presynaptic structural-functional analysis provides direct support for the following mechanistic steps: (1) LGI1 autoantibodies interfere with LGI1’s endogenous function of increasing the presynaptic potassium channels density. (2) The reduction of presynaptic potassium channels prevents efficient repolarization of the presynaptic action potential. And (3) the resulting longer presynaptic action potential increases release probability.

### Comparison with LGI1 knock-out mice

Consistent with increased release probability upon antibody application, knock-out of LGI1 in mice increased transmission at hippocampal CA3-CA3 synapses^17^ and in CA1 neurons^22,55^. Furthermore, overexpression of LGI1 decreased synaptic strength at perforant-path granule cell synapses.^20^ The synaptic strengthening upon LGI1 knock-out is probably mediated presynaptically by an increased release probability, because LGI1 knock-out postsynaptically either decreased AMPAR clustering and quantal size^12,23,25,27^ or did not affected quantal size ^20,22^. However, some synapses show no presynaptic effect upon LGI1 knock-out or LGI1 application. For example, in CA1 neurons PPR was mostly unaffected by LGI1 application^12^ or LGI1 knock-out^12,23,25,55^. These differences in the presynaptic effect of LGI1 knock-out on synaptic transmission may relate to the differential expression of LGI1, with highest expression in the hippocampal outer and middle molecular layers of the denate gyrus (perforant path– granule cell synapses).^18^ Furthermore, LGI1-overexpression shortened presynaptic action potentials in primary hippocampal cultures, leading to lower action potential-evoked calcium entry and hence glutamate release.^56^ Therefore, LGI1 autoantibodies induce effects that are reminiscent of those observed in LGI1 knock-out mice and thus support the mechanistic model that LGI1 increases the presynaptic potassium channel density, shortens the presynaptic action potential duration, lowers the release probability, and thereby dampens neuronal activity.

### Presynaptic functions of LGI1 outside of the release site

We found that K_v_1 channels are homogeneously distributed across the axon and bouton and only a small minority of potassium channels was located at the release site (Fig. 4C). Furthermore, LGI1 autoantibodies decreased the K_v_1 density within and outside of the Bassoon-labeled release sites, indicating a homogeneous reduction throughout the bouton (Fig 5). Our data therefore argue that LGI1, in addition to its trans-synaptic alignment, controls potassium channels also outside of the release site. Indeed, it was recently shown that LGI1 autoantibodies also alter the K_v_1 cluster distribution at the axon initial segment.^57,58^ The autoantibody-induced increase in neuronal excitability was mediated by antibodies specifically targeting the LRR-domain.^30,31,57,58^ Consistently, structural analyses indicate that LGI1 can form protein complexes in a cis-configuration serving as an extracellular scaffold instead of a transsynaptic hub.^15,59^ It remains to be determined if the density of presynaptic K_v_1 channel outside of the release site is controlled by LGI1 proteins in the cis-configuration.

### Calcium channels do not mediate the increased release probability

The analyses of calcium channels were motivated by the increase in synaptic release probability following LGI1 autoantibody treatment, a phenomenon typically observed upon changes in calcium channel density or function. Furthermore, proteome data previously indicated an interaction of the LGI1-receptor ADAM22 with pore-forming calcium channel alpha-subunits and their beta-subunits^19,38^, which control calcium channel surface expression and kinetics.^41,60^ We used STED and EM imaging of Ca_v_2.1 calcium channels and direct electrophysiological recordings of presynaptic calcium current density and gating kinetics. Yet, we found neither presynaptic Ca_v_2.1 channel abundance nor calcium current amplitude and channel gating kinetics were strongly affected by LGI1 autoantibodies (if anything, there was a reduction in the channel density). Therefore, potential effects of LGI1 on presynaptic calcium channels do not contribute to the increased release probability induced by LGI1 autoantibodies.

### Monoclonal autoantibodies targeting the EPTP-domain of LGI1 lead to enhanced AP broadening

We used previously characterized patient-derived monoclonal autoantibodies to specifically target the EPTP or the LRR domain of LGI1.^31^ While EPTP-targeting autoantibodies led to enhanced broadening during train stimulation in our recordings, LRR autoantibodies did not affect action potential broadening. The EPTP-domain of LGI1 has previously been shown to mediate binding to ADAM22, and EPTP-targeting autoantibodies hence prevented binding of LGI1 to ADAM22.^13,27,30,31^ Both EPTP and LRR autoantibodies reduced K_v_1.1 protein levels in hippocampus-enriched solubilized brain lysates but the effect seemed stronger with EPTP-compared to LRR autoantibodies.^31^ In contrast, some studies observed an increased neuronal excitability only with LRR but not with EPTP autoantibodies.^31,57,58^ Our data indicate that the duration of the action potential is preferentially affected by EPTP autoantibodies, whereas LRR-targeting antibodies were previously shown to interfere with multimerization and cause internalization of the LGI1-ADAM22 complex.^13,30,31^ A differential effect of autoantibodies targeting EPTP and LRR is conceivable because of the complex interplay of various types of potassium channels in controlling excitability and action potential repolarization.^61^ However, more studies are needed to understand the differential effect of the subunit-specific autoantibodies on excitability and action potential repolarization. Furthermore, although we tested two monoclonal antibodies for each LGI1 functional domain, our data cannot rule out that LRR autoantibodies with different binding epitopes other than those tested here are able to affect presynaptic K_v_1 function. These interesting mechanistic differences in the regulation of neuronal and synaptic function by the two domains of LGI1 may also be reflected in differential symptoms associated with mutations in these domains, such as auditory features that occur less frequently in congenital epilepsy caused by EPTP truncation compared to LRR truncation ^62^.

## Acknowledgements

We thank Claudia Sommer for expert technical assistance, the Electron Microscopy Facility of IST-Austria for resources, and Tereza Belinova in the Imaging and Optics Facility of IST-Austria for 3D reconstruction.

## Supplementary material

Supplementary material is available online.

