## Supplementary material for "LGI1 autoantibodies enhance synaptic transmission by presynaptic K_v_1 loss and increased action potential broadening"

**for**

### eFigures and Figure legends

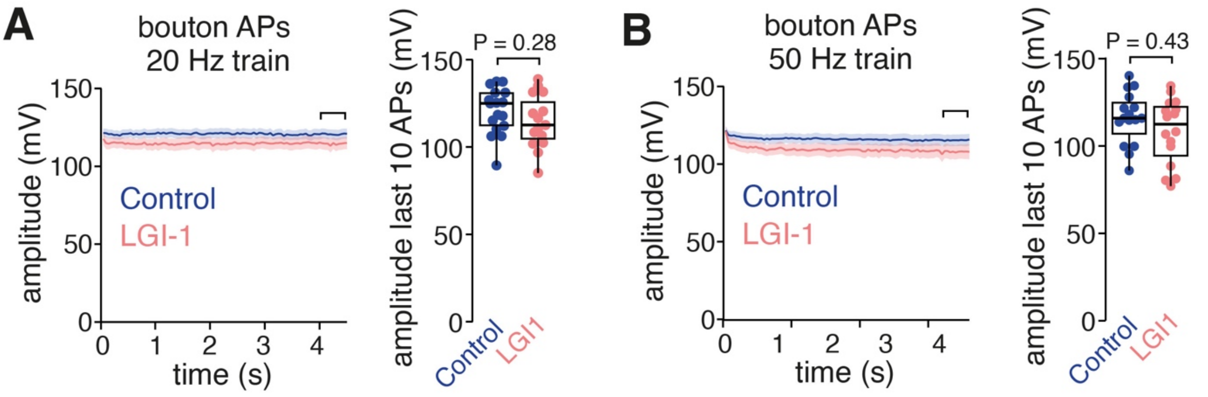


**eFigure 1 LGI1 autoantibodies do not affect bouton action potential amplitude**

**(A)** Time course (left) and magnitude of action potential amplitude change (right; mean amplitude of 10 last train action potentials) during 20 Hz train stimulation in control and LGI1 autoantibody-treated presynaptic boutons (blue and orange, respectively; amplitude normalized to that of the first train action potential).

**(B)** Bouton action potential amplitudes as in A for 50 Hz train stimulation.

Box plots provide median and cover percentile 25–75, whiskers reflect percentiles 10–90. Numbers in brackets provide number of recorded presynaptic boutons or somata.

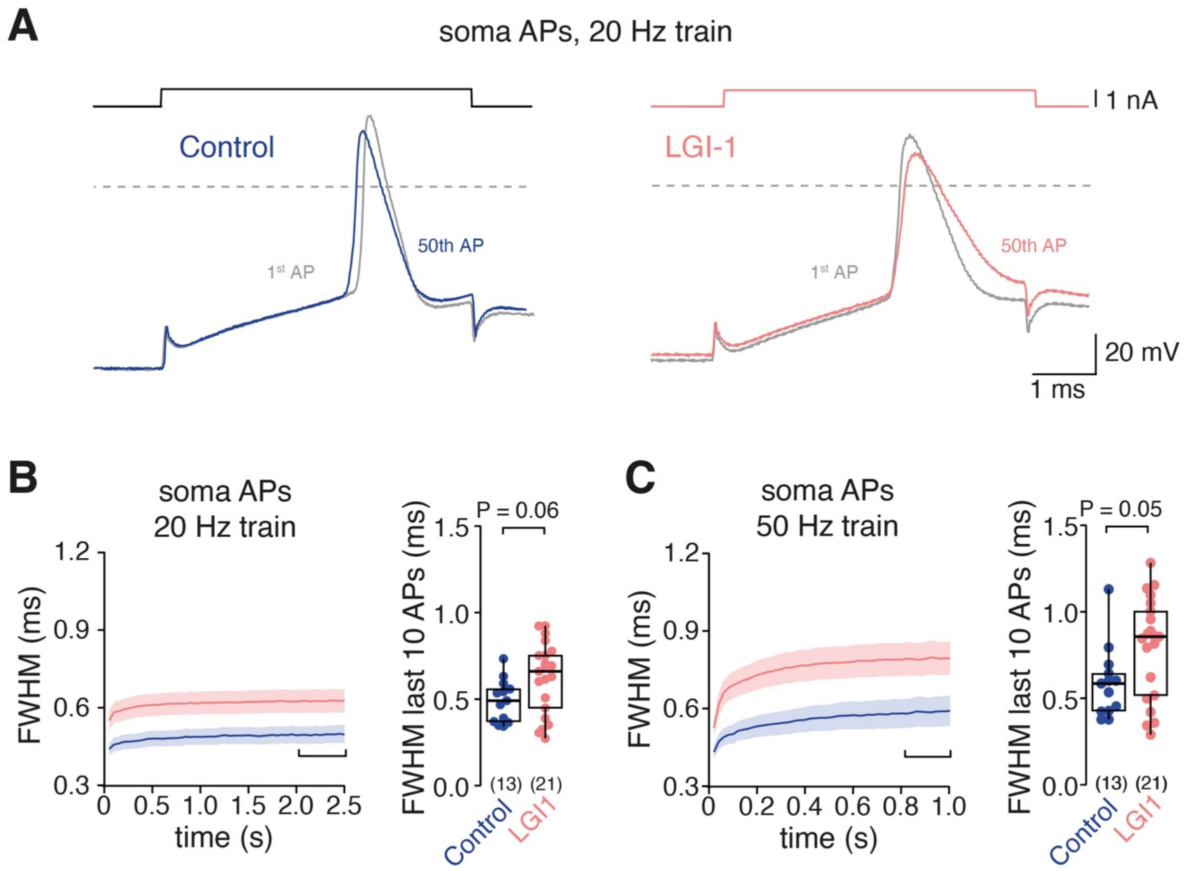


**eFigure 2 LGI1 autoantibodies cause somatic action potential broadening**

- **(A)** Superposition of the 1st (grey) and 50th somatic action potential during 20 Hz train stimulation under control condition (blue) and following LGI1 autoantibody-treatment (orange).
- **(B)** Time course (left) and magnitude of somatic action potential broadening (right; mean FWHM of 10 last train action potentials) during 20 Hz train stimulation in control and LGI1 autoantibody-treated cells (color code as in A; FWHM normalized to that of the first train action potential).
- **(C)** Soma action potential broadening as in B for 50 Hz train stimulation.

Box plots provide median and cover percentile 25–75, whiskers reflect percentiles 10–90. Numbers in brackets provide number of recorded presynaptic boutons or somata.

### eMethods

#### Animals

All experimental and breeding animals were kept under a 12 h light and 12 h dark cycle with water and food *ad libitum*. Electrical recordings and STED imaging were performed on primary cultures of Sprague Dawley (SPRD) rat neurons, SIM imaging was performed on primary cultures of C57Bl/6J mouse neurons. Electron microscopy was performed on hippocampal brain tissue of adult C57Bl/6J mice after intraventricular passive transfer of human polyclonal IgG fractions (n = 9 mice for control and LGI1 antibody infusion each). Surgery and implantation of intraventricular catheters and osmotic pumps was performed as described previously.[1,2] Brain tissue was harvested after 14 days of continuous antibody infusion. Animals were handled according to the regulations of the Federal Saxonian, Thuringian (licence # UKJ-17-053), and Bavarian state authorities (license # 55.2.2-2532-2-811) and in accordance with European regulations (Directive 2010/63/EU).

#### Human LGI1 antibodies

LGI1 IgG was purified from plasma exchange material of 3 patients suffering from LGI1 encephalitis and high-titer (1:100) serum anti-LGI1 autoantibodies and of control patients without CNS disorders and without antineuronal antibodies according to previously published protocols.[3] All patients provided informed consent for use of plasma exchange material and use of human material was approved by the local ethics committee of Jena University Hospital (licence # 2019-1415-Material). [4]For treatment with monoclonal domain-specific antibodies, recordings with two antibody clones targeting the same domain were merged (LRR-mAb02, LRR-mAb03, EPTP-mAb12 and EPTP-mAb04)[4] and are referred to as EPTP or LRR autoantibodies.

#### Antibody incubation of neuronal cultures

Purified polyclonal serum autoantibodies from patients with LGI1 encephalitis and patient-derived monoclonal EPTP or LRR autoantibodies were pre-diluted in 100 µl culture medium and added to cells grown in 1 ml culture medium seven days before recordings at final concentrations of 100 µg/ml or 10 µg/ml for polyclonal and monoclonal autoantibodies, respectively. One day before recordings, half of the medium (550 µl) was removed from wells and purified polyclonal or monoclonal autoantibodies were added to the remaining medium (final concentration 100 µg/ml or 10 µg/ml, respectively). For one-day incubation (LGI1-1d), autoantibodies were applied similar one day before recordings only. Purified control antibodies from patients without LGI1-encephalitis and monoclonal control antibodies were applied identical to the respective autoantibody groups. For controls for train EPSC recordings (Fig. 1), control antibodies were applied one day before recordings.

**Neuron cell culture for electrical recordings and STED imaging**

Neuronal cultures were generated from postnatal day (P)0 - 2 SPRD rat pups. For individual cultures, both hippocampi from only one individual pup were used. Following decapitation, hippocampi were isolated on ice-cold L15 medium and cut into 1 mm-sized chunks. Chunks were digested in Trypsin/EDTA (calcium- and magnesium-free) for 5 minutes at 37° C, dissociated in ice-cold serum medium (DMEM + 5% FKS) using fire-polished Pasteur-pipettes of decreasing diameter, passed through a 70 µm cell strainer and centrifuged for 8 min at 100g at 4° C. The pellet was then resuspended in serum medium and cells were counted before plating 50.000 cells on PDL-coated (50 kDa; 1:50 suspension) 13-mm coverslips. Cells were incubated for 48h (at 37° C under steam-saturated carbogen atmosphere) in serum medium before the full medium was replaced by Neurobasal A-medium containing B27 (1:50), GlutaMax (1:100), Gentamicin (1:1000), FKS (final 1%), and AraC (4 µM). From 4 days after plating, medium was exchanged by the same Neurobasal A-based medium without AraC and cells were grown until antibody-application.

**Neuron cell culture for SIM imaging**

Primary neurons were prepared from E18 C57Bl / 6J mice. After isolation of single cells, from hippocampal tissue using TrypLE, 100000 neurons were cultivated/plated on 18 mm high precision coverslips (Carl Roth, LH23.1). The coverslips were pretreated with 25% HCl and 99% EtOH and coated with 300 µl (0.1 mg/ml) poly-D-lysine (PDL, Sigma-Aldrich) for 1 hr at 37° C. Neurons were cultured in 1.6 ml of Neurobasal containing 1% Glutamax, 2% B27 Plus supplement (all purchased from Life Technologies), and 5 µg/ml gentamycin (Sigma Aldrich), and 50% medium was replaced weekly. Cultures were fixed at DIV14 using 4% methanol (MeOH)-free paraformaldehyde (PFA, Life Technologies) for 10 minutes at room temperature (RT) and then stored in 1x PBS at 4°C.

**ExM-reagents**

Acrylamide (AAm; catalog no. A4058), N,N'methylenebis(acrylamide) (BIS; catalog no. M1533) and sodiumacrylate (SA; catalog no. 408220), ammonium persulfate (APS; catalog no. A3678) and N,N,N',N'-tetramethylethylenediamine (TEMED; catalog no. T7024) were purchased from Sigma-Aldrich. APS and TEMED were diluted to 10% (w/v) stock solutions in double-destilled $H_2O$ and purchased from Sigma-Aldrich. Sodium Dodecyl Sulfate (SDS; catalog no.AM9820) was purchased from Invitrogen and sodium chloride (NaCl; catalog no. S5886) was purchased from Sigma-Aldrich. Tris- (hydroxymethyl)-aminomethan (Tris; catalog no. 1.08382) was purchased from Sigma-Aldrich.

**Neuron fixation and permeabilization**

Fixed primary neurons were rinsed three times with 1x PBS and permeabilized with 0.2% Triton$^{TM}$ X-100 (ThermoFisher; catalog no. 28314) in 1x PBS. The neurons were then washed three times with 1x PBS and crosslinked overnight applying 4% FA + 30% AAm (Sigma; catalog no. A4058) in 1x PBS at 37°C.

**Gelation, denaturation and antibody labeling and post gelation immunostaining**

Samples were gelled under nitrogen supply with monomer solutions containing 1.1M SA + 2.0M AA + 90ppm / 0.009% BIS + 1x PBS. Polymerization was initiated by addition of 1.5 ppt TEMED and 1.5 ppt APS. The gelation chamber and the monomer solution were cooled to 4°C. Gels were incubated for 1.5 h at RT in a nitrogen-filled and humidified chamber. Next, the gels were detached from the 1 mm coverslips and transferred to a 5 min preheated 95°C denaturation solution (200mM SDS + 200mM NaCl + 50mM Tris) and incubated for 1h at 95°C. After the samples cooled to RT for 10 min, they were transferred into a Petri dish containing Millipore water for 15 min in the first expansion step. Water was replaced with 1x PBS three times to shrink the gel to its initial size for immunolabeling. Samples were incubated overnight (~15 h) with monoclonal anti-Bassoon antibody (Enzo Life Sciences; catalog no. SAP7F407), polyclonal anti-Vglut1 (Synaptic Systems; catalog no. 135304),polyclonal anti-Kv1.1 (Frontier Institute; catalog no. AB_2571787), or polyclonal anti-Kv1.2 (Frontier

Institute; catalog no. AB_2571789) diluted 1:100 in 5% bovine serum albumin (BSA; Sigma; catalog no. A7030 in 1x PBS). Incubation of primary antibodies was performed on a spinning wheel at RT. Gels were then washed four times with 0.1% PBST (Tween 20; Sigma; catalog no. P9416 in 1x PBS) for 20 min on a spinning wheel at RT. Samples were subsequently incubated with commercially available AF488 anti-guinea pig antibodies (Jackson Immuno Research, catalog no. 706545148), CF568 anti-rabbit antibodies (Sigma; catalog no. SAB4600400) and custom-conjugated ATTO643 anti-mouse antibodies (ATTO TEC; catalog no. AD643-31; DOL: 1.9; protein concentration: 0,57 g/l) for 3 h applying a 1:100 dilution in in 5% BSA. The secondary antibody incubations were performed applying constant agitation on a heating block at 37°C. Following immunostaining samples were washed four times with 0.1% PBST for 20 min on a spinning wheel at RT. The hydrogels were expanded in double-distilled $H_2O$ until they reached their maximum size by exchanging $H_2O$ in 30 min intervals.

**ExM sample mounting**

After expansion, the gels were mounted on poly-D-lysin-coated (Sigma; catalog no.: P6407; diluted 1:10 in Millipore water) one-well chambers with 0.17 mm thick glasses (Cellvis; catalog no.: C1-1.5H-N).

**Image acquisition and processing**

All Lattice SIM$^2$ images were acquired on a ZEISS Elyra 7 equipped with HR Diode 488 nm – 500 mW (green), HR DPSS 561 nm – 500 mW (red) and HR Diode 642 nm – 500 mW (far red) lasers in excitation pathway. The images were acquired by using the LBF 405/488/561/642 laser blocking filter sets and SBS LP 560 beam splitter and emission was collected on sCMOS PCO Edge 4.2M cameras. The images were obtained by using a C-Apochromat 63x1.15 NA water objective. The ZEISS ZEN 3.0 SR FP2 (black) software (ZEISS) was used to control imaging parameters. All Z-stacks were collected with fast-frame mode and 91 nm slice interval. All images were SIM$^2$ processed by using the ZEISS ZEN 3.0 SR FP2 (black) software. (It's a two-step reconstruction algorithm: First, order combination, denoising and median filtering were performed). Subsequently, iterative deconvolution is applied. SIM$^2$-processed images were visualized by using FIJI/ImageJ software. Brightness and contrast of all SIM$^2$-images was adjusted linearly. The obtained pixel size of the SIM images is 31 nm.

**Colocalization analysis**

Analysis of CF568 ($K_v$1.1/2) and ATTO643 (Bassoon) colocalization within the same pixel size was conducted on 22 optical slices (2.1 µm total axial range, slice thickness of 91 nm within each z-stack). The extent of within-pixel fluorescent signal colocalization was quantified using the "colocalization threshold" plugin and measured in ImageJ (Version 2.3.0). For each channel a threshold is determined automatically to avoid subjective bias. Colocalization was calculated in each optical slice and then collapsed into a maximal z-projection to reveal colocalized pixels across the entire image thickness.

**Freeze-fracture electron microscopy**

Freeze-fracture replica labeling was performed as previously described [5,6]. Briefly, mice were perfused with 2% paraformaldehyde, 15% saturated picric acid in 0.1M phosphate buffer (PB), pH 7.4, for 13 min at rate 7 ml/min. Coronal hippocampal slices of 120 µm thickness were prepared with Leica Vibratome VT1000s (0.30 mm/sec speed at 75 Hz frequency) in 0.1 M PB kept at ice-cold temperature. Slices were infiltrated with 30% glycerol in Tris-buffered saline, pH 7.4 (TBS) overnight at 4°C. Slices were trimmed (1.2 mm diameter including the CA1 area to the dentate gyrus in the dorsal hippocampus) and sandwiched between a pair of copper carriers with 70 µm thick double-sided tape and then high pressure frozen (Bal-tec HP010). Frozen samples were stored in liquid nitrogen until use. Mirrored carbon-platinum-carbon replicas (5-2-25 nm thickness for each) were prepared using either Bal-tec BAF060 or Leica ACE900 freeze fracture machines. Tissue replica was solubilized with 2.5% SDS in 20% sucrose-TBS (SDS buffer) for 18 hours at 80°C, 50 rpm continuous shaking. Replicas were washed in ceramic well plates with fresh SDS buffer followed by 3 x wash buffer (0.1% BSA in TBS) for 20 min at room temperature (RT). Non-specific binding sites were blocked 20 min with blocking solution (3% bovine serum albumin (BSA), 2% cold water fish skin gelatin and 0.05% Tween20 in TBS) 1 hour at RT and then replicas were incubated in guinea pig anti-$Ca_v$2.1 antibody (Frontier Institute Cat# VDCCa1A-GP, RRID:AB_2571851) diluted to 4 µg/µl in antibody diluent, overnight at 15°C. After washing in wash buffer (0.1% BSA in TBS) for 20 min three times, replicas were incubated in goat anti-guinea pig IgG conjugated with 5 nm colloidal gold (BBI solutions) diluted 1:50 overnight in humid chamber at 15°C. Following washing in wash buffer (3 times) and TBS for 20 min at RT each, replicas were washed in MilliQ water 6 times and picked up onto formvar coated TEM grids and air-dried. Replicas were examined in FEI Technai12 transmission electron microscope at 120 kV acceleration voltage. Representative >20 images from presynaptic active zones of either the performant path-granule cell or mossy fiber-CA3 pyramidal cell synapses in the dentate gyrus were taken at 52,000x magnification on Veleta CCD camera with Radius software (EMSIS). Active zone areas were indicated with the aggregation of intramembrane particles in shallow dents on the replica as described previously,[5,6] and were delineated using Darea software.[7] Analysis of active zone area and numbers of gold particles labeling $Ca_v2.1$ were assessed using Darea software to determine $Ca_v2.1$ density.[7]

**Pre-embedding electron microscopy**

Pre-embedding immunolabeling was adapted from methods previously described [8,9]. Mice were perfused with 4% paraformaldehyde, 15% saturated picric acid in 0.1M phosphate buffer, 0.05% glutaraldehyde, pH 7.4, 13 min at rate 7ml/min. Coronal hippocampal sections of 80 µm thickness were prepared with Leica Vibratome VT1000s (0.30 mm/sec speed at 75 Hz frequency) in 0.1 M PB kept at ice-cold temperature. Sections were infiltrated with increasing concentrations up to 30% sucrose in 0.1M PB overnight at 4°C before two cycles of permeabilization by snap freezing in liquid nitrogen and rapid thawing on 50°C hotplate for 2 min. Sections were washed 3 times in TBS and then free aldehyde groups quenched with 50mM glycine in TBS for 10 min at RT. Non-specific binding sites were blocked for 20 min with blocking solution. Then, sections were incubated with primary antibodies diluted to 4 µg/µl; either $K_v1.1$ (rabbit polyclonal antibody, Frontier Institute, Cat.#: MSFR103600, RRID: AB_2571787) or $K_v1.2$ (rabbit polyclonal antibody, Frontier Institute, Cat.#: MSFR103650, RRID: AB_2571789) diluted in antibody diluent (AD; 1% BSA, 1% CWFSG and 0.05% Tween20 in TBS). Sections were incubated in Pelco Biowave® Pro+ Microwave, 150 W at 25°C with alternating cycles each 5 min (on, off) without vacuum for 2 hrs. Sections were washed in washing buffer (0.1% BSA in TBS) for 3 min (150 W, 1 min alternating on, off cycles), three times, without vacuum. Then sections were incubated with secondary antibody (Nanoprobes #2004, Fab' goat anti-rabbit IgG 1.4nm gold) diluted 1:200 in AD solution overnight at 4°C. Next day, sections were washed in TBS (three times) and PBS (twice) for 3 min cycles in microwave (150 W, 1 min alternating on, off cycles). Sections were post-fixed in 1% glutaraldehyde in 0.1M PBS for 40 sec in microwave (100 W) followed by treatment with glycine 50 mM in PBS for 40 sec in microwave (100 W). After PBS washing and MilliQ rinsing (40 s 100 W, three times each), sections were treated with silver amplification (Nanoprobes HQ silver enhancement kit, #2012) for 9 min at RT. Then, sections were washed in MilliQ three times followed by osmification (0.5% $OsO_4$ in MilliQ) for 15 mins at RT and another three washes in MilliQ (40 s 100 W). After incubation with 1% uranyl acetate solution for 30 mins in dark at RT, sections were washed in MilliQ for 40 sec, 150 W MW followed by dehydration in ethanol series (30%, 50%, 70%, 90%, 96% and 2x 100%), each step at 40 sec, 100 W MW. An intermediate solution of propylene oxide, 2x 40 s (MW 250 W) was followed by 50:50 propylene oxide to Durcopan epoxy resin (Durcopan ACM, Science Services GmbH) infiltration, 3 min (250 W, 1 min alternating on, off cycles) under vacuum (20 mmHg; 30 sec alternating on, off cycle). Similarly, infiltration with freshly prepared pure Durcopan resin 250 W MW with vacuum infiltration, 2x 3 min was followed by flat embedding within ACLAR film sandwich and polymerization at 60°C overnight. Next day, region of interest was trimmed with a scalpel and re-embedded into BEEM capsule with fresh Durcopan resin. Ultrathin sections of 70nm thickness were cut using Leica UC7 ultramicrotome with Diatome® Diamond knife. Ribbons of 25-30 serial sections were applied to each formvar-coated slot TEM grids. Resin sections on grids were contrasted with 1% uranyl acetate 15 min and 2% Reynold's lead citrate 7 min, before examination in FEI Technai12 transmission electron microscope at 120 kV acceleration voltage. Images from dentate gyrus mossy fibers at 52,000x magnification were recorded using Radius imaging software with Veleta CCD camera (EMSIS). Axonal series spanning 19 to 24 sections were aligned and virtually stacked using Fiji-ImageJ, and multiple axon terminals were tracked to unmyelinated axons through 7-10 multiple images. Silver-enhanced gold particles for $K_v1.1$ or $K_v1.2$ were counted for individual axon series and the ratios of particles in presynaptic active zone, perisynaptic sites (within 60 nm from active zone) and extrasynaptic sites were calculated. In addition, representative reconstructed $K_v1.1$ axons were transformed to a 3D model using IMARIS software (Fig. 4D).

**Immunofluorescence stainings of $K_v1.1$, $K_v1.2$, and $Ca_v2.1$**

Immunocytochemcial stainings of potassium channels in primary cultures we performed as previously described.[10] In short, the cells were fixed with 4% paraformaldehyde (PFA) in PBS at room temperature (RT) for 12 min. Following a washing step with PBS to remove residual

PFA from the samples, the cells were permeabilized with PBS containing 0.3% Triton-X 100 at RT for 3 min at 37°C. Afterwards, the cover slips were blocked three times for 15 min in a solution containing PBS, 10% fetal calf serum (FCS), 25 mM glycine, and 2% bovine serum albumin (BSA). Then the cells were incubated overnight with either anti-$K_v$1.1 (rabbit polyclonal antibody, 1:400, Frontier Institute, Cat.#: AB_2571787), anti-$K_v$1.2 (rabbit polyclonal antibody, 1:400, Frontier Institute, Cat.#: AB_2571789), or anti-$Ca_v$2.1 (rabbit polyclonal antibody, 1:1000, Synaptic Systems, Cat.#: 152203) together with anti-Bassoon (mouse monoclonal antibody, 1:500, Enzo Life Sciences, Cat.#: SAP7F407), and anti-VGluT1 (guinea pig polyclonal antibody, 1:1000, Synaptic Systems, Cat.#: 135 304) in blocking solution at 4°C. The bound antibodies were visualized by corresponding fluorophore conjugated secondary antibodies for 1.5 h at RT. Finally, all samples were stained with DAPI for 30 min at RT and embedded afterwards in Mowiol.

**STED image acquisition and analysis**

We used an Expert Line Abberior microscope system with an Olympus (UPlanXApo) 60x oil objective with a numerical aperture of 1.42. The system contained 3 pulsed fluorescence excitation laser modules for wavelengths of 485, 561, and 640 nm, the corresponding filter cubes (GFP, Cy3, Cy5), and avalanche photodiode (APD) detectors. For STED imaging the system uses a pulsed high-power STED laser module with a wavelength of 775 nm (3 W laser power, repetition rate of 40 MHz). Each image was obtained with sequential line scanning by switching between the needed excitation lasers line-wise during each recording. Each image contained a three channel-recording for either $K_v$1.1, $K_v$1.2 or $Ca_v$2.1 channels, the active zone protein Bassoon and vGluT1. All channels were imaged with a dwell time of 1 µs/pixel and a tenfold line accumulation. Based on the secondary antibody signals, the excitation laser power for each channel was adjusted to prevent over- or underexposure effects. Excitation laser power for $K_v$1.1, $K_v$1.2 or $Ca_v$2.1 (secondary antibody STAR Red, laser module 640 nm) was set to 2.61% with a STED laser power of 35% and for Bassoon (secondary antibody STAR Orange/CF568, laser module 561 nm) to 5.26% with a STED laser power of 10%. For vGluT1 (secondary antibody Alexa Fluor 488, laser module 488 nm) only confocal images were acquired with an excitation laser power of 4%. The settings were kept constant throughout each set of experiments.

A custom-written macro of the software Fiji[11] was used to generate a mask for either Bassoon or vGlut1 based on a prior adjusted threshold. The average pixel intensity of $K_v1.1$, $K_v1.2$ or $Ca_v2.1$ was quantified for each spot within either of the Bassoon or vGlut1 masks.

**Postsynaptic EPSC recordings**

Coverslips from treated cultures were mounted on an upright microscope (Nikon FN1), visualized using difference-interference optics with an 60x water-immersion objective (Nikon; NA 1.0) and perfused with ACSF containing (in mM): 125 NaCl, 3 KCl, 25 Glc, 25 $NaHCO_3$, 1.25 $Na_2HPO_4$, 2 $CaCl_2$, 1 $MgCl_2$, pH 7.4. Excitatory AMPA-receptor postsynaptic currents (EPSCs) were pharmacologically isolated by including 20 μM AP-5 and 10 μM gabazine in the ACSF. To reduce recurrent or run-away excitation, EPSC amplitudes were reduced by including 2 mM of kynurenic acid into the bath solution. EPSC train recordings were performed at room temperature (22–24°C) and as previously described [12,13]. In short, somatic whole-cell voltage-clamp recordings were performed using pipettes pulled from borosilicate glass (Science Products, Hofheim, Germany; DMZ Universal Electrode Puller; Zeitz Instruments, Martinsried, Germany) to resistances between 5 and 8 MOhm and filled with internal potassium gluconate-based solution containing (in mM): 150 K-gluconate, 3 Mg-ATP, 0.3 Na-GTP, 10 K-HEPES, 10 NaCl and 0.05 EGTA, 3 QX-314 chloride, pH adjusted to 7.2 by KOH, osmolarity 293 mOsm. A second external glass electrode filled with ACSF was used to evoke EPSC trains by short 0.1 ms external stimulations 40 times at 20 or 50 Hz. Only those experiments in which stimulation evoked rapidly-rising, monophasic EPSCs with a single peak and constant initial and steady-state train amplitudes were considered for analysis. Currents were recorded with an EPC10/2 amplifier (HEKA Elektronik, Lambrecht/Pfalz, Germany). Postsynaptic EPSCs were filtered with the internal 3 kHz 4-pole Bessel filter and sampled at 20 kHz (EPSC trains) or 200 kHz (PPRs). EPSC amplitudes were analyzed as the phasic component as described previously.[12] The paired-pulse ratio (PPR) was calculated as the ratio of the second to the first train EPSC amplitude. Data were analyzed using Igor Pro, version 6.32A (WaveMetrics, Lake Oswego, OR, USA), extended by Patcher's Power Tool (http://www3.mpibpc.mpg.de/groups/neher/index.php?page=aboutppt, version 2.19) and the NeuroMatic plug-in[14] (http://www.neuromatic.thinkrandom.com/; version 2.00) as well as self-written analysis routines.

### Presynaptic action potential recordings

Presynaptic current-clamp action potential recordings were performed as described previously.[15,16] In brief, quartz glass pipettes were pulled from Quartz glass pipettes (without filament; Heraeus Quartzglas, Kleinostheim, Germany) by a DMZ Universal Electrode Puller with oxygen-hydrogen burner (Zeitz Instruments, Martinsried, Germany)[17] to resistances of 6–13 MΩ. Pipettes were filled with a potassium gluconate–based internal solution (composition see *Postsynaptic EPSC recordings* section) and membrane potentials were corrected for the calculated 12 mV liquid junction potential offline. Recordings were performed at physiological temperature (34.5–36.0°C) using a Multiclamp 700A patch-clamp amplifier (Molecular Devices, San Jose, USA). Pipettes were mounted on a micromanipulator (Kleindiek Nanotechnik, Reutlingen, Germany) by a custom-build pipette holder and pipette capacitance was systematically minimized (for details on capacitance reduction and compensation see[15]). All current-clamp recordings were filtered with the internal 10 kHz 4-pole Bessel filter of the Multiclamp 700A amplifier and digitized (200 kHz) with the HEKA EPC10/2 using Patchmaster software (HEKA Elektronik, Lambrecht/Pfalz, Germany). Holding currents were adjusted to hold membrane potentials at –65 mV (not liquid-junction potential corrected) and bridge compensation was adjusted to minimize the current injection artifacts. Action potentials were analyzed using Igor Pro, version 6.32A (WaveMetrics, Lake Oswego, OR, USA) with the Patcher's Power Tool (http://www3.mpibpc.mpg.de/groups/neher/index.php?page=aboutppt; version 2.19) and NeuroMatic plug-in[14] (http://www.neuromatic.thinkrandom.com/; version 2.00) as well as self-written analysis routines. Action potential trains were evoked by 0.3-ms current injections (90 times) at 20 and 50 Hz. Current-injection amplitudes were adjusted such that action potentials were initiated at or after the end of current-injection, thereby reducing the impact of current-injection artifacts on recorded voltages. Action potential amplitudes were quantified between the membrane potential preceding current-injection and action potential peak. Action potential half-durations were quantified at the voltage of half-maximal amplitude.

### Somatic action potential recordings

Somatic current-clamp action potential recordings were performed as described previously.[18] In brief, neurons cultured for 17 to 21 days *in vitro* were perfused with ACSF containing (in mM): 138 NaCl, 20 Glucose, 10 HEPES, 2.5 KCl, 1.2 $Mg_2SO_4$, and 2 $CaCl_2$, pH 7.4 at room temperature. Electrophysiological recordings were conducted by a HEKA EPC-10 amplifier

(HEKA Elektronik, Lambrecht/Pfalz, Germany) with a sampling rate of 20-100 kHz and low-pass filtered at 2.9 kHz using the amplifier's Bessel filters. Somatic whole-cell current-clamp recordings were performed using pipettes pulled from borosilicate glass (Science Products, Hofheim, Germany; DMZ Universal Electrode Puller; Zeitz Instruments, Martinsried, Germany) to resistances between 3 and 5 MOhm and filled with internal potassium gluconate-based solution containing (in mM): 150 K-gluconate, 10 HEPES, 10 NaCl, 3 MgATP, 0.3 $Na_2GTP$, and 0.05 EGTA, pH adjusted to 7.3 by KOH, osmolarity 290 mOsm.

**Presynaptic calcium current recordings**

Calcium currents were pharmacologically isolated in an external recording solution containing (in mM): 120 NaCl, 10 HEPES, 4 TEA-Cl, 1 4-AP, 2.5 KCl, 10 Glucose, 1.1 $MgCl_2$, 1.1 $CaCl_2$, 0.01 TTX. Recording electrodes were filled with an internal recording solution containing (in mM): 90 $CsMeSO_4$, 10 NaCl, 10 HEPES, 20 TEA-Cl, 20 $K_2$-Phosphocreatine, 4 MgATP, 0.3 NaGTP, 0.1 ETGA, pH adjusted to 7.35 using CsOH. For bouton identification, 60–200 µM Atto 488-Carboxy (ATTO TEC, Siegen, Germany) were added to the internal solution. Calcium currents were recorded at physiological temperature (34.5–36°C) in voltage-clamp mode and corrected for leak and capacitive currents using the P/4-method. For activation, 3-ms step depolarizations (10 mV steps from –80 to +20 mV) were applied and average current amplitudes in the last millisecond of the test pulse were analyzed at different depolarized potentials. Only experiments with rapid and complete inactivation after the end of the depolarization step were analyzed. The time course of current activation within the first millisecond after pulse onset was fit with an exponential function with delayed onset and a time constant ($\tau_m$): $I(t) = I_\infty(1 - \exp(-(t - t_0)/\tau_m))$. For deactivation of calcium currents, 10-ms steps from 0 mV to voltages between −10 and −80 mV were applied and current amplitudes during the last millisecond of the inactivation step were analyzed. For inactivation time constant, single exponentials were fit to the first 3 milliseconds of inactivation steps manually adjusted to start following the peak of the calcium tail current. Calcium currents were filtered with the internal 10 kHz 6-pole Bessel filter and sampled at 200 kHz using a HEKA EPC10/2 amplifier (HEKA Elektronik, Lambrecht/Pfalz, Germany).

**Statistics**

All data are reported as median [IQR], if not stated otherwise. The Shapiro-Wilk test was used to test for normal distribution of the data points and a significance level of 0.05 was used. For the multiple comparisons (Fig. 1 or 7), non-parametric ANOVA (Kruskal-Wallis) tests were performed followed by non-parametric post-hoc tests (Dwass-Steel-Critchlow-Fligner pairwise comparisons) or Dunn's multiple comparison tests. The calculations were performed with jamovi[19] or OriginPro 2019.

**Data availability**

Data that support the findings of this study are available from the corresponding author on reasonable request.

7. Kleindienst D, Montanaro J, Bhandari P, Case MJ, Fukazawa Y, Shigemoto R. Deep Learning-Assisted High-Throughput Analysis of Freeze-Fracture Replica Images

Applied to Glutamate Receptors and Calcium Channels at Hippocampal Synapses. *Int J Mol Sci*. 2020;21(18):1-19. doi:10.3390/IJMS21186737
